# Subphase Material Stabilizes Films of Pulmonary Surfactant

**DOI:** 10.1101/2020.02.27.968099

**Authors:** K. Andreev, M. W. Martynowycz, I. Kuzmenko, W. Bu, S. B. Hall, D. Gidalevitz

## Abstract

When compressed by the shrinking alveolar surface area during exhalation, films of pulmonary surfactant *in situ* reduce surface tension to levels, at which surfactant monolayers collapse from the surface *in vitro*. Vesicles of pulmonary surfactant added below these monolayers slow collapse. X-ray scattering here determined the structural changes that improve stability. Grazing incidence X-ray diffraction on monolayers of extracted calf surfactant detected an ordered phase. Mixtures of dipalmitoyl phosphatidylcholine and cholesterol, but not the phospholipid alone, mimic that structure. At concentrations that stabilize the monolayers, vesicles in the subphase had no effect on the unit cell, and the film remained monomolecular. The added vesicles, however, produced a concentration-dependent increase in the diffracted intensity. These results suggest that the enhanced resistance to collapse results from components of an ordered interfacial phase which partition from subphase to the surface, increasing the area of the ordered structure.

**SIGNIFICANCE:** Low alveolar surface tensions are essential for maintaining the integrity of the pulmonary air-sacks during normal breathing. Films of pulmonary surfactant cause the low tensions. The interfacial structures required for the low surface tensions remain uncertain. These studies used X-ray scattering to determine the initial structure of pulmonary surfactant monolayers, and to establish how vesicles of pulmonary surfactant enhance the ability of those initial monolayers to sustain low tensions. The initial monolayers contained ordered structures that differ from the crystalline forms widely speculated to occur in alveolar films. The added vesicles had no effect on the local structure of the initial monolayer, but substantially increased the area of the ordered regions. This structural change reasonably explains the functional improvement.

## INTRODUCTION

Pulmonary surfactant is the combination of lipids and proteins that lowers surface tension (γ) in the lungs (1). The material forms a thin film on the surface of the liquid that lines the alveolar air-sacks (2). When compressed by the decreasing interfacial area during exhalation, the films reduce γ to exceptionally low values, below 5 mN·m^−1^ (3–8). These low γ indicate that the films avoid a phase transition. Surfactants adsorb to an air-water interface only until they reach the equilibrium spreading tension (γ_e_, ~25 mN·m^−1^ for fluid-phase phospholipids) (9), defined by coexistence of the two-dimensional monolayer with its three-dimensional bulk phase (10). For fluid films, attempts to increase the interfacial density further, either by adding constituents or decreasing area, produce at most a transient decrease in γ before constituents collapse from the interface to restore equilibrium.

The γ levels in the lungs, well below γ_e_, indicate that the alveolar film avoids this transition, and that it must be metastable (11).

The structure of the alveolar film that sustains low γ remains unclear (12). Vesicles of pulmonary surfactant adsorb as collective packets (13–15), delivering their complete set of constituents to form initial monolayers at the interface (16). At physiological temperatures, such monomolecular films, whether formed by adsorption or by spreading from solutions in volatile solvents, readily collapse at the γ_e_ (11, 17). Additional vesicles, however, added below these initial monolayers, substantially increase their stability at low γ (17, 18). The studies reported here focus on the structural changes that produce that improved stability.

Two models propose structures that could explain the stabilizing effect of the additional material. The first, or classical model (19–21), contends that the film at low γ is tilted-condensed (TC). The alkyl chains of phospholipids in that structure form a two-dimensional crystalline array (22). TC films readily undergo compression to low γ. At 37°C, dipalmitoyl phosphatidylcholine (DPPC), which is the most prevalent constituent of pulmonary surfactant (~35%) from most species (23), forms the TC phase above γ_e_ (24, 25). Monomolecular films with the complete biological mixture of surfactant phospholipids form a disordered structure surrounding TC domains, which contain essentially pure DPPC (26). Vesicles in the subphase could provide a source of additional DPPC that might enrich the interfacial content of that compound and enlarge the TC domains to dominate the film (21).

A second model proposes that the additional material stabilizes the initial monolayer by increasing its thickness (27–29). Electron microscopy has shown that at least portions of the alveolar film contain multiple layers (30–32). If those additional layers form during adsorption, they might impede collapse and enable the films to sustain low γ. Material in the subphase would achieve its stabilizing effects by producing a more rigid, multilamellar film.

We used X-ray scattering from liquid surfaces to investigate the divergent structural hypotheses of the two models. The classical model predicts that pulmonary surfactant added to the subphase should increase the area occupied by a TC phase. Diffraction of X-rays incident on the surface at a grazing angle should detect the distinct signal of the TC structures. Material in the subphase should increase that signal. The multilamellar model predicts instead that the added material should form additional layers, detectable by X-ray reflectivity (XRR). Our experiments provide structural information on the initial monolayers formed by extracted calf surfactant, and how additional surfactant added below the surface changes those structures.

## MATERIALS AND METHODS

### Materials

CLSE was obtained from ONY, Inc. CLSE is prepared by centrifugal pelleting of large lipoprotein aggregates lavaged from freshly excised calf lungs (33), followed by extraction of the particles with nonpolar solvent (34). For preparation of surfactant vesicles, evaporation of the solvent under a stream of nitrogen followed by an overnight incubation in vacuum yielded the dried lipid-protein mixtures. CLSE preparations were then resuspended in HEPES-buffered saline (HS: 10 mM 4-(2-hydroxyethyl)-1-piperazineethanesulfonic acid pH 7.0, 150 mM NaCl) at a final phospholipid concentration of 32 mM (~100 mg·mL^−1^) followed by three cycles of sequential freezing and thawing (–80/23 °C), with vigorous vortexing before each freezing. Further sonication for 30 min in a water-bath with ice yielded the final dispersion. Dynamic light scattering (ZetaPALS™, Brookhaven Instruments Corporation, Nova Instruments LLC, Holtsville, NY) determined the size and polydispersity of freshly prepared liposomes to characterize the dispersions. DPPC and cholesterol were purchased from Avanti Polar Lipids (Alabaster, AL), and used without further purification or characterization.

### Formation of interfacial films

Our experiments used the air-water interface in a Langmuir trough filled with HEPES-buffered saline containing calcium (HSC: 10 mM HEPES pH 7.0, 150 mM NaCl, 1.5 mM CaCl_2_) to mimic the surface of the liquid layer in an alveolus. Aliquots of CLSE or lipids dissolved in chloroform (Sigma-Aldrich, St. Louis, MO) were spread onto the aqueous surface until γ fell slightly below 70 mN·m^−1^. The films were then compressed using a Teflon barrier while monitoring γ with a Wilhelmy plate. GIXD was measured at regular intervals. Vesicles of CLSE, dispersed in HS, were either deposited as small droplets at the interface, or injected into the subphase to allow interfacial adsorption. Deposition of droplets containing dispersed CLSE continued until γ reached ~26 mN·m^−1^. For adsorbed CLSE, the final concentrations were 0.75 or 1.50 mM phospholipid.

### Surface X-ray scattering

X-ray measurements were conducted at either beamline 9-ID-C or 15-ID-C of the Advanced Photon Source at Argonne National Laboratory (Argonne, IL). The experiments used a Langmuir trough mounted in a hermetically sealed, helium-filled canister. The oxygen level, monitored continuously, was <1% to minimize background scattering. Temperature was the ambient level of 23°C for all experiments.

#### GIXD

GIXD provides information about the in-plane, lateral order of films. X-rays incident at a grazing angle penetrate only ~100 Å below the surface, and are therefore sensitive to the Langmuir monolayer rather than the bulk subphase. Measurements of GIXD used an X-ray beam with a wavelength (*λ*) of 0.9202 Å (9-ID-C) or 1.24 Å (15-ID-B), striking the surface at an incident angle corresponding to 0.85% of the critical value (*q*_c_ = 0.0217 Å^−1^). GIXD measurements record the diffracted intensities expressed as a function of the wave-vector parallel to the interface, *q_xy_*. For an X-ray beam below the critical angle, intensity is a function of the momentum transfer parallel to the interface, *q_xy_*. At an angle of 2θ, *q*_xy_ is given by:

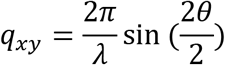

The scattered intensity can be described in terms of diffraction from a large number of randomly oriented, two-dimensional crystalline domains. After subtracting linear background, the intensities were analyzed in OriginPro software (version 8.0, Northampton, MA) by fitting the data to a single Lorentz-Gauss (1:1) crossed peak. The *q*_xy_ of the peak yielded the repeat distances between the diffracting hydrocarbon chains:

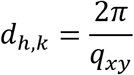

The breadth of the peak provided the coherence length, *L*_xy_, of the diffracting region according to the Sherrer formula (35),

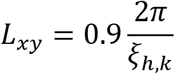

where *ξ* = (FWHM^2^ – Δ^2^)^1/2^ for a full width of the peak at the half-maximum (FWHM), and Δ is the resolution of the Soller slits. The acceptance of the Soller slits, fixed at 1.4 milliradians, determined errors in *q*_xy_ (9.56 × 10^−3^ Å^−1^).

The nearest neighbor tilt in our films follows the equation:

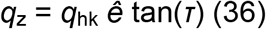

where *τ* is the angle of molecular tilt along the **a** dimension of the unit cell (37, 38), indicated by *ê* (39).

#### Specular XRR

XRR determines the electron density profile across the interface. Measurements record the reflected intensities, expressed as a function of the wave-vector perpendicular to the interface, *q*_z_. For a beam incident on the surface at an angle *α*, *q*_z_ is given by:

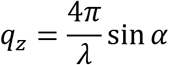

The measured intensity of the reflected beam, corrected for off-specular background, is normalized relative to the incident intensity. The beam’s footprint limited measurements of XRR to the range of *q*_z_ from 0.01 to 0.60 Å^−1^. For determinations of the electron density profile, solutions to the phase problem used a model-independent approach. The interface was modeled as ~50 discrete slabs of continuous electron density and length of 1/3 Å. These slabs were allowed to vary in order to match the measured reflectivities and maintain a smooth curve in the program StochFit (40). This continuous electron density was then fit to a simplified model of two slabs of discrete, uniform density (*ρ*) and length (*l*), bounded by the known densities of the subphase (water) and superphase (air). This approach has been recently applied for studying membrane interaction mechanisms by antimicrobial peptide mimics (41, 42–44), lipopolysaccharide permeability in *Salmonella spp*. (45, 46), and surface catalysis (47).

## RESULTS

### Initial monolayers of CLSE

We first determined the structure of compositionally well-defined monomolecular films containing the full set of constituents in extracted calf surfactant (calf lung surfactant extract, CLSE). Deposition of CLSE in solutions of a volatile solvent formed monolayers at low interfacial densities. These Langmuir films have served as a model of the initial alveolar film since the earliest research on pulmonary surfactant (48). Grazing incidence X-ray diffraction (GIXD) provided structural information.

When compressed, diffraction first occurred at γ = 55 mN·m^−1^. The intensities were best fit by two broad distributions centered at *q*_xy_ = 1.36 and 1.47 Å^−1^ (Fig. 1 *A*). This pattern corresponded to a centered rectangular unit cell containing two acyl chains (37), with dimensions of *a* = 8.54 Å and *b* = 5.45 Å, and an area per alkyl chain of 23.3 Å^2^. Diffraction was centered at *q_z_* = 0 Å^−1^, both here and at other γ, indicating the absence of molecular tilt between nearest neighbor alkyl chains. Further compression to γ = 40-45 mN·m^−1^ increased the intensities and slightly reduced the size of the unit cell, but preserved the space group. At γ = 26 mN·m^−1^, the pattern had changed (Fig. 1 *A*). A single peak at *q*_xy_ = 1.49 Å^−1^ indicated conversion to a hexagonal unit cell (37), where *a* = *b* = 4.87 Å with an area per unit cell of 20.6 Å^2^ (Table 1).

**FIGURE 1.**
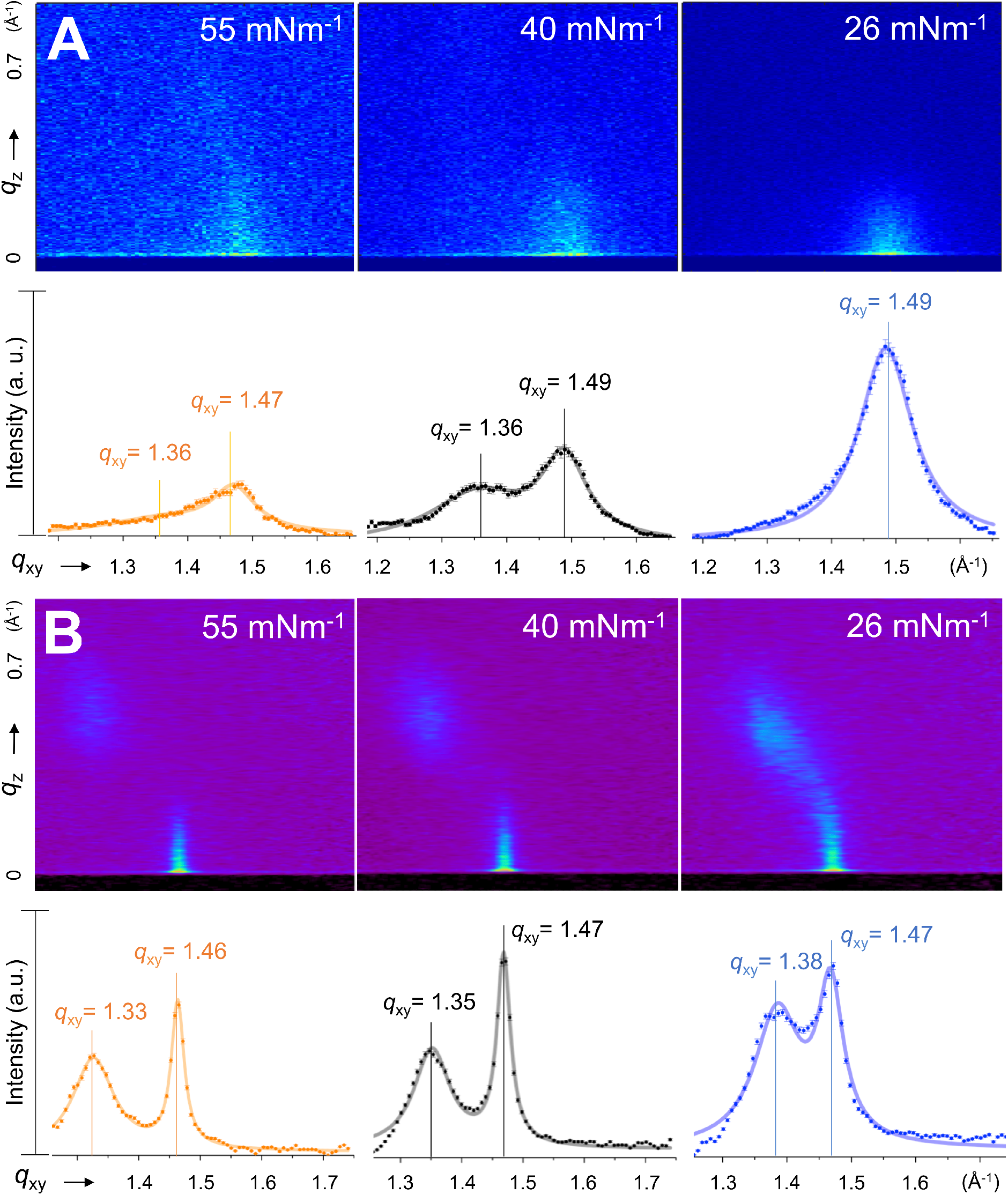
GIXD from monolayers of (*A*) CLSE and (*B*) DPPC spread from organic solvent and compressed to different γ. The upper row gives the imaged intensities. The lower row provides the variation of intensities integrated over *q*_z._ Continuous curves represent the best fits to the data using Lorentz-Gauss crossed peaks. Errors are assumed to be Poisson-distributed.

**Table 1.**
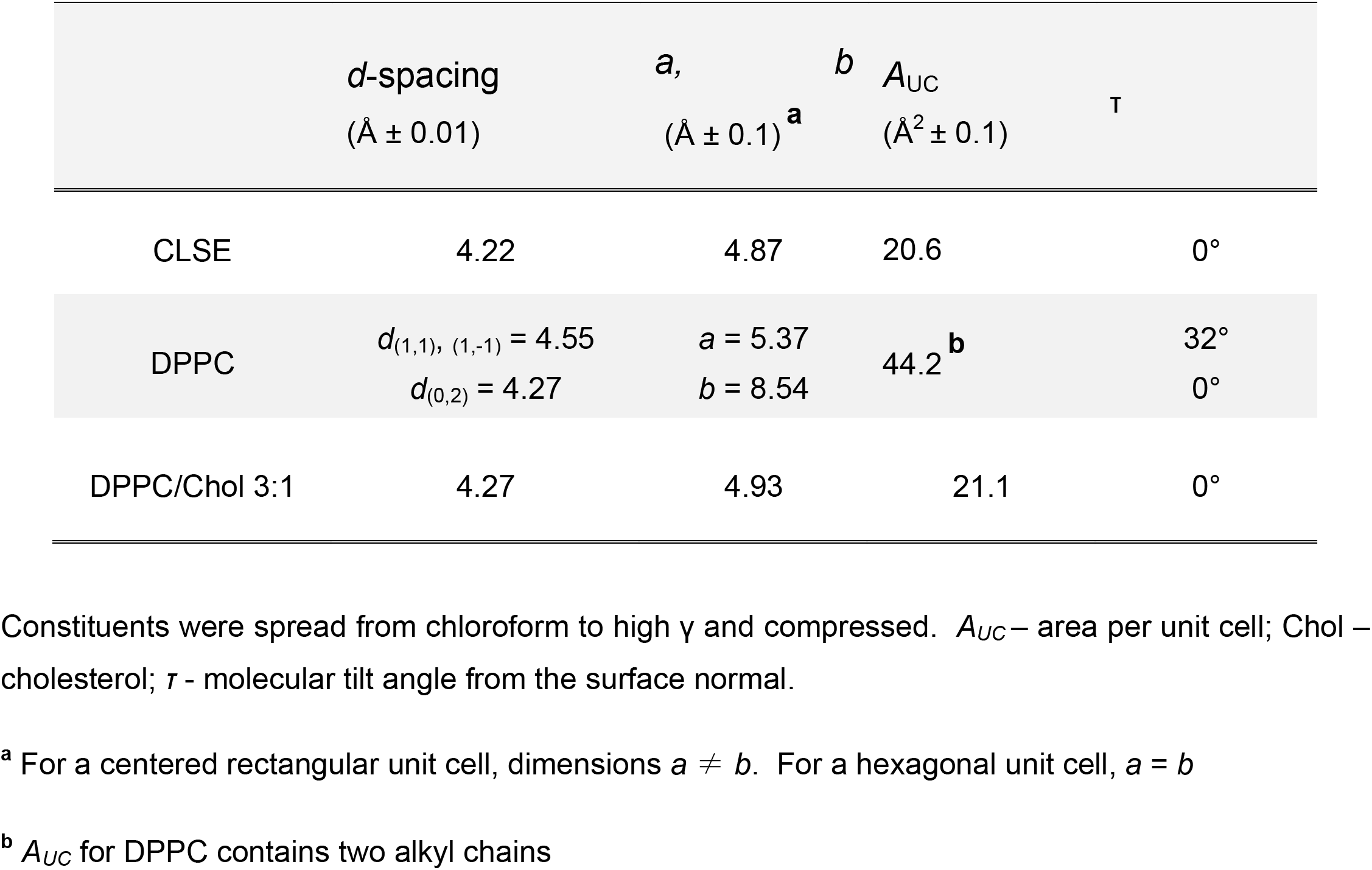
Lateral organization of crystalline regions (γ = 26-28 mN·m^−1^)

| | *d*-spacing<br>($\text{\AA} \pm 0.01$) | *a*,<br>($\text{\AA} \pm 0.1$)<sup>a</sup> | *b*<br>*A*<sub>UC</sub><br>($\text{\AA}^2 \pm 0.1$) | $\tau$ |
| --- | --- | --- | --- | --- |
| CLSE | 4.22 | 4.87 | 20.6 | 0° |
| DPPC | $d_{(1,1), (1,-1)} = 4.55$<br>$d_{(0,2)} = 4.27$ | $a = 5.37$<br>$b = 8.54$ | 44.2<sup>b</sup> | 32°<br>0° |
| DPPC/Chol 3:1 | 4.27 | 4.93 | 21.1 | 0° |

Constituents were spread from chloroform to high $\gamma$ and compressed. *A*<sub>UC</sub> – area per unit cell; Chol – cholesterol; $\tau$ - molecular tilt angle from the surface normal.
<sup>a</sup> For a centered rectangular unit cell, dimensions $a \neq b$ . For a hexagonal unit cell, $a = b$
<sup>b</sup> *A*<sub>UC</sub> for DPPC contains two alkyl chains

According to the classical model, the ordered regions that resist collapse should be composed of TC DPPC. The diffraction from CLSE, however, at γ just above γ_e_, where collapse begins, differed significantly from previously reported results for DPPC (49). To confirm the pattern produced by DPPC under our experimental conditions, we measured GIXD from monolayers containing only that compound. Monomolecular films at 55 mN·m^−1^ produced the expected two pronounced peaks at *q_xy_* ≈ 1.33 Å^−1^ and 1.46 Å^−1^ (Fig. 1 *B*), corresponding to a centered rectangular unit cell with dimensions of *a* = 5.2 Å, *b* = 8.5 Å, and an area of 44.2 Å^2^, or 22.1 Å^2^ per alkyl chain. In contrast to the pattern for CLSE, the shift off plane of the second peak (*q_z_* ≈ 0.7 Å^−1^) indicated tilt of acyl chains towards nearest neighbors. Compression to γ = 26 mN·m^−1^ produced a partial merger of the two peaks, suggesting a decrease in the *d*-spacing of the (1,1) and (1,−1) planes from 4.72 to 4.55 Å (Table 1). The molecular tilt angle also decreased. A transition to a hexagonal unit cell did not occur. The diffraction patterns for ordered regions of CLSE and TC DPPC indicated different structures, and suggested distinct compositions.

Prior studies have suggested that ordered domains in monolayers of CLSE contain cholesterol as well as DPPC (26, 50). Measurements of GIXD determined if cholesterol/DPPC mixtures could mimic the structure of ordered regions in CLSE (49). At γ ≈ γ_e_, GIXD from DPPC with 25 mol% cholesterol closely approximated the signal from CLSE. A single diffraction peak at *q*_xy_ = 1.47 Å and *q*_z_ ≈ 0 Å^−1^ indicated alkyl chains hexagonally packed with a unit cell of *a* = *b* = 4.93 Å and *A_UC_* = 21.1 Å^2^, without detectable molecular tilt (Table 1). The results supported the prior finding that, in addition to DPPC, the ordered regions of CLSE contained significant cholesterol.

### Effect of subphase CLSE

To determine how vesicles in the subphase affected the structure of the interfacial film, we first established whether monolayers formed by aqueous dispersions and solutions in chloroform had the same structure. CLSE dispersed in aqueous buffer and deposited at the surface produced the same GIXD as the Langmuir films. Diffraction again showed the single peak (Fig. 2 *A*) centered at *q_z_* = 0, indicating a hexagonal lattice without molecular tilt. The dimensions (*a* = 4.97 Å) and *d*-spacing (4.30 Å) were slightly larger than the values for CLSE spread from organic solvent (Table 2), but the structures were basically comparable.

**Table 2.** Effect of subphase material on adsorbed CLSE films (γ = 26 mN·m^−1^)

| CLSE subphase<br>concentration (mM)<sup>a</sup> | *d*-spacing<br>($\text{\AA} \pm 0.01$) | *a*,<br>($\text{\AA} \pm 0.1$) | *b*<br>*A*<sub>UC</sub><br>($\text{\AA}^2 \pm 0.1$) | $L_{xy}$<br>($\text{\AA} \pm 2$) | Total Intensity<br>(a.u.) |
| --- | --- | --- | --- | --- | --- |
| 0.00 | 4.30 | 4.97 | 21.4 | 65 | 207±14 |
| 0.75 | 4.30 | 4.97 | 21.4 | 58 | 693±26 |
| 1.50 | 4.27 | 4.93 | 21.1 | 87 | 868±29 |

$L_{xy}$ – in-plane coherence length
<sup>a</sup> Phospholipid concentration

**FIGURE 2.**
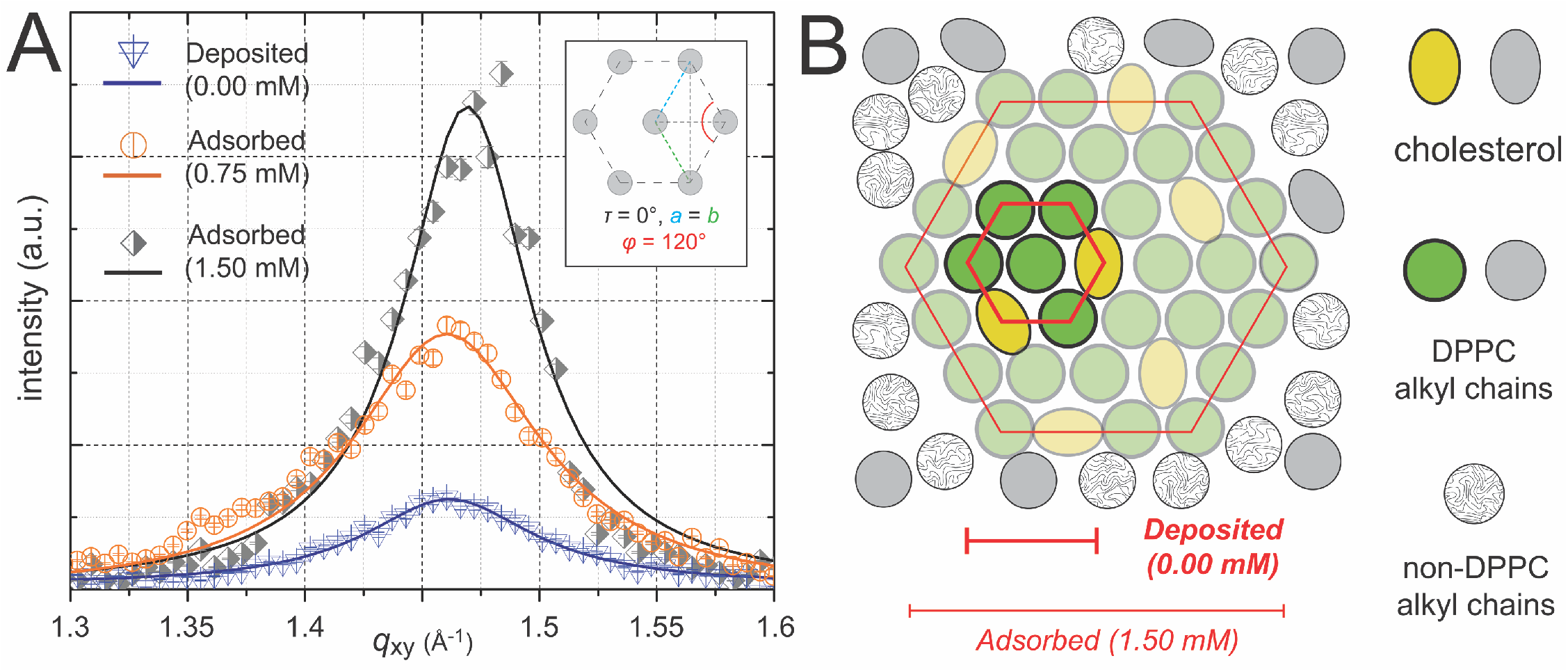
Effect of subphase material on the diffracted intensity. (*A*) GIXD from CLSE films at γ = 26 mN·m^−1^ with additional surfactant below the surface at the indicated phospholipid concentrations. Symbols indicate experimental measurements with error assumed to have Poisson distributions. Continuous curves give the best fit of Lorentz-Gauss peak-functions to the data after subtraction of background. Fits were weighted by their statistical significance. The diagram in the insert represents the unit cell of the hexagonal lattice. *a, b,* φ – parameters of the unit cell; *τ* - angle of molecular tilt from the surface normal. (*B*) Cartoon illustrating a possible arrangement of lipids within the crystalline domains (colored) and in the surrounding disordered phase (gray).

Vesicles added to the subphase produced minimal change in *q_xy_* of the single peak (Fig. 2 *A*), with no effect on *q_z_*. The ordered regions retained a hexagonal unit cell with comparable dimensions and untilted chains (Table 2). The total diffracted intensity, however, increased. With a concentration of 0.75 mM, the integrated intensity was more than threefold higher than the signal for the monolayer above an empty subphase (Table 2). The effect was concentration-dependent (Fig. 2 *A*). Doubling the subphase-concentration to 1.50 mM increased the intensity further by 25% (Table 2). During exposure to an incident beam of equal intensity for equivalent durations, the area of an ordered structure should determine the diffracted intensity. Although material in the subphase had no effect on the molecular arrangement within ordered regions, the vesicles significantly increased their total area.

The presence of 1.50 mM CLSE in the subphase also decreased the width of the diffracted peak. The coherence length, *L*_xy_, which indicates the average size of individual ordered domains, is inversely related to the width (see Materials and methods). The additional material increased *L*_xy_ by 50% (Fig. 2 *B*, Table 2). Our studies provided no information concerning the number of ordered regions. They offered no insight into whether subphase vesicles caused nucleation of new ordered domains. Our results did show that the increased ordered area occurred at least in part by growth of the existing domains.

### Transverse structure of adsorbed CLSE

X-ray reflectivity provided the electron density along the normal to the interface (Fig. 3). A simplified model of the CLSE films as a stack of two horizontal, homogenous slabs successfully fit all measured intensities (Fig. 3 *A*). The electron densities for the upper, nonpolar slab 1 varied from 0.314 to 0.317 e^−^·Å^−3^ (0.94−0.95 relative to water, *ρ*_aqua_) (Fig. 3 *B*). These slightly exceeded the densities of 0.304 e^−^·Å^−3^ and 0.307−0.311 e^−^·Å^−3^ for pure DPPC (51), and dipalmitoyl phosphatidylglycerol (DPPG), respectively (41, 52). In turn, the densities of the polar slab 2 ranging within 0.428-0.431 e^−^·Å^−3^ (1.26-1.29·*ρ*_aqua_) were lower than those for phospholipids (Table 3) (41, 51–54). These differences seemed most likely to reflect the presence of cholesterol and the surfactant proteins in CLSE. Material added to the subphase had little or no effect on the density or thickness for either slab. The subphase vesicles therefore produced essentially no change in the total thickness of the films. For all measurements, the thickness of 26-27 Å (Table 3) indicated the presence of a monolayer. Like the lateral structure detected by GIXD, material in the subphase had no detectable effect on the transverse structure.

**FIGURE 3.**
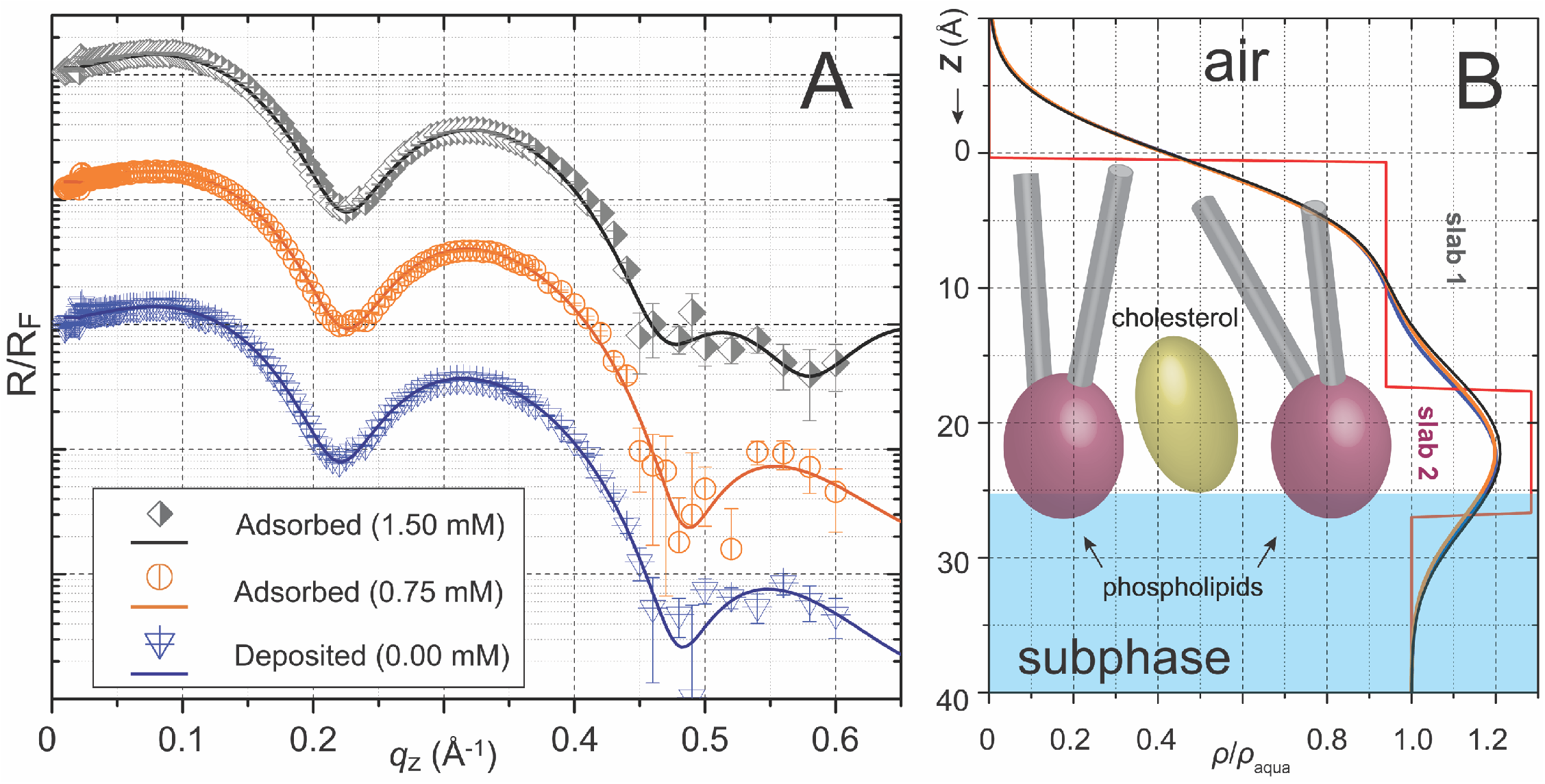
Effect of subphase material on the transverse structure. (*A*) X-ray reflectivity data from films of CLSE at γ = 26 mN·m^−1^. The reflected intensity, *R,* is normalized relative to the Fresnel reflectivity, *R*_F_, for an ideally flat air-water interface. Symbols give measured values, where vertical bars indicate the error assuming Poisson distributions about the counted intensity. The continuous curves give the best fit to the data by the Fourier transform of the two-slab model of electron density. (*B*) Electron density profiles, normalized relative to the electron density of aqueous buffer (*ρ*_aqua_ ~ 0.334 e^−^·Å^−3^), derived from the data in panel *A*. *Z* denotes the distance from the top of the upper slab (slab 1). The molecular cartoon suggests the general orientation of lipids in the monolayer.

**Table 3.** Two-slab model of CLSE films perpendicular to the interface

| CLSE subphase concentration (mM) | Slab 1<br>(non-polar) |  | Slab 2<br>(polar) |  |
| --- | --- | --- | --- | --- |
| | $l_n$ (Å) | $\rho_n / \rho_{\text{aqua}}$ | $l_p$ (Å) | $\rho_p / \rho_{\text{aqua}}$ |
| 0.00 | 17.0 ±0.07 | 0.94 ±0.01 | 9.40 ±0.15 | 1.28 ±0.01 |
| 0.75 | 16.7 ±0.09 | 0.95 ±0.01 | 9.14 ±0.17 | 1.29 ±0.01 |
| 1.50 | 15.8 ±0.09 | 0.94 ±0.02 | 11.15 ±0.17 | 1.26 ±0.01 |

$l$ and $\rho$ are the thickness and electron density, respectively, of the nonpolar (subscript n) and polar (subscript p) homogeneous slabs. Values of $\rho$ are normalized relative to the aqueous buffer ( $\rho_{\text{aqua}} \sim 0.334 \text{ e}^- \cdot \text{\AA}^{-3}$ ). Errors correspond to model uncertainty.

## DISCUSSION

Our results address a fundamental question concerning the function of pulmonary surfactant. Monomolecular films of CLSE *in vitro* fail to reproduce its behavior *in situ*. In excised lungs, CLSE fully replicates the performance of endogenous surfactant (55, 56). *In vitro*, however, at physiological temperatures, monomolecular films of the same material collapse at γ_e_ (11). A series of studies over several years have suggested that vesicles injected below the surface stabilize the films (17, 18, 57). At the same concentrations that slow collapse by two orders of magnitude (17), our experiments show structural changes. These results provide significant insights into how the alveolar film becomes capable of sustaining very low γ.

Our results show that the additional material has no effect on the local structure of the initial monolayer. The thickness of the film remains monomolecular. Prior speculation that formation of a multilamellar film accounted for the greater stability appears unfounded. The local lateral structure, although different from the prediction of the classical model, also remains unaltered. The additional material, however, produces a major increase in the ordered area of the film. Both the growth of the ordered regions and their lateral structure have functional significance.

### Growth of the ordered phase

Our findings provide new insights into a major problem with the classical model. TC monolayers replicate the performance of the alveolar film, but the model fails to explain how that crystalline film would form. DPPC represents 35-40% of the surfactant phospholipids from most species (58). In monolayers of the phospholipids from CLSE, the TC phase that coexists with fluid regions contains pure DPPC (59). Formation of a TC film would require a major compositional change of the initial monolayer, and enrichment of DPPC.

Earlier versions of the model have invoked enrichment by selective exclusion. Compression below γ_e_ of coexisting solid and fluid phases causes collapse of the fluid regions. If compressed far enough, the result is a solid film (19–21). At physiological temperatures, however, and γ just above γ_e_, ordered domains in films of CLSE occupy ~4% of the interface (60). During a physiological compression, exclusion of fluid regions sufficient to produce a solid film seems unlikely.

Our results support instead a compositional change that occurs by selective insertion. Material added to the subphase would provide a source of additional DPPC. Preferential partitioning of DPPC into the interfacial monolayer, perhaps facilitated by the surfactant proteins, would enrich the film in that compound. An ordered phase composed of DPPC with cholesterol could then grow. The process is dose-dependent. The concentration of the alveolar liquid is unknown, but likely to be above 100 mM phospholipid (57), much higher that the values used here. The extent of ordered regions in the alveolar film should be correspondingly greater.

The basis of partitioning would be thermodynamic rather than kinetic, but not from a difference in γ_e_. Phospholipids above their main melting transition share a common γ_e_ of ~24 mN·m^−1^ (9, 61). Phospholipids in solid structures adsorb poorly, suggesting that γ_e_ may instead be just below the γ for a clean interface (9, 62). Insertion of compounds according to these γ_e_ would enrich the film in compounds that form fluid structures rather than DPPC. We speculate that constituents instead partition based on spontaneous curvature. DPPC, with its cylindrical shape, might prefer the relatively flat air-water interface over the curvature of undulating structures.

### Structure of the ordered phase

Vesicles injected below the surface, which slow collapse (17), and increase the ordered area (Fig. 2), have no discernible effect on the local structure. The ordered regions retain a hexagonal unit cell. The dimensions of the lattice are unchanged, and the acyl chains remain untilted. The film remains monomolecular. No differences in the local structure can explain the greater stabilities at low γ.

The ordered regions in monolayers of CLSE are not simply DPPC in the TC phase. The diffracted intensity from that structure generates the dual peaks of the centered rectangular lattice with off-axis signal from tilted chains. The films of CLSE instead produce the single peak of a hexagonal lattice with the equatorial intensity of untilted chains. The TC phase that is the cardinal feature of the classical model is absent.

Prior microscopic studies (26, 50), and our results here suggest instead that the ordered regions in CLSE are liquid-ordered (L_o_). Sufficient cholesterol converts TC DPPC to L_o_ structures (49). The phospholipid accommodates the cholesterol in an ordered structure that diffracts, but the space group shifts, and the acyl chains lose their tilt (49). The ordered regions in CLSE have these characteristics, and closely resemble the structure of DPPC with 25% cholesterol. The prior microscopic studies yielded parallel results. Monolayers of the phospholipids in CLSE form coexisting phases (59). The ordered domains contain only DPPC (59), and have the characteristics of the TC phase. Physiological levels of cholesterol convert the fixed, irregular domains in the phospholipid films to circular shapes that can reconfigure rapidly (26). The solid, TC domains change to the fluid characteristics of the L_o_ phase (63). The prior and current studies, using different experimental approaches (64), both conclude that the cholesterol in CLSE converts TC domains to L_o_ (65).

The lack of apparent change in local structure by the added material poses an important functional question. Growth of the ordered phase would stabilize the compressed film only if that structure resists collapse. The L_o_ phase may lack that feature. Collapse of a two-dimensional film into the third dimension at γ_e_ indicates the ability to flow that defines a fluid (66). Domains of the L_o_ phase act like a two-dimensional fluid. They lack long-range order (67), and undergo rapid reconfiguration (26). In sufficient amounts, cholesterol induces films of DPPC to collapse (68–70). We speculate that the cholesterol in the ordered regions of CLSE may be insufficient to allow collapse.

## CONCLUSIONS

X-ray scattering from monolayers of pulmonary surfactant detects an ordered phase. DPPC with cholesterol, but not DPPC alone, mimics those ordered regions. Surfactant vesicles added below the surfactant film has no effect on the local structure, including its thickness and the unit cell of the ordered regions. The subphase material, however, substantially increases the ordered area in a concentration-dependent manner.

## AUTHOR CONTRIBUTIONS

KA, MWM, SBH and DG designed the research. KA, MWM, IK and WB performed X-ray scattering experiments. MWM, KA, SBH, and DG analyzed experimental data. KA, MWM, SBH, and DG wrote the paper.

## ACKNOWLEDGMENTS

The authors thank Dr. Edmund Egan (ONY Inc., Amherst, NY) for the gift of CLSE. This research was supported by funds from the National Institutes of Health (HL130130 and 136734). This study used resources of the Advanced Photon Source, a U.S. Department of Energy (DOE) Office of Science User Facility operated for the DOE Office of Science by Argonne National Laboratory under Contract No. DE-AC02-06CH11357. The illustrations for Cover Art were kindly provided by Marta Kosacheva.

## REFERENCES

1. Goerke, J., and J. A. Clements. 1985. Alveolar surface tension and lung surfactant. Handbook of Physiology - The Respiratory System. Vol. III, Part 1. P. T. Macklem and J. Mead, editors. American Physiological Society, Washington, D.C., pp. 247–261.

2. Bastacky, J., C. Y. Lee, J. Goerke, H. Koushafar, D. Yager, L. Kenaga, T. P. Speed, Y. Chen, and J. A. Clements. 1995. Alveolar lining layer is thin and continuous: low-temperature scanning electron microscopy of rat lung. J. Appl. Physiol. 79(5):1615–1628.

3. Fisher, M. J., M. F. Wilson, and K. C. Weber. 1970. Determination of alveolar surface area and tension from in situ pressure-volume data. Respir. Physiol. 10:159–171.

4. Horie, T., and J. Hildebrandt. 1971. Dynamic compliance, limit cycles, and static equilibria of excised cat lung. J. Appl. Physiol. 31(3):423–430.

5. Schürch, S., J. Goerke, and J. A. Clements. 1976. Direct determination of surface tension in the lung. Proc. Natl. Acad. Sci. USA. 73(12):4698–4702.

6. Valberg, P. A., and J. D. Brain. 1977. Lung surface tension and air space dimensions from multiple pressure-volume curves. J. Appl. Physiol. 43(4):730–738.

7. Wilson, T. A. 1981. Relations among recoil pressure, surface area, and surface tension in the lung. J. Appl. Physiol. 50(5):921–930.

8. Smith, J. C., and D. Stamenovic. 1986. Surface forces in lungs. I. Alveolar surface tension-lung volume relationships. J. Appl. Physiol. 60(4):1341–1350.

9. Lee, S., D. H. Kim, and D. Needham. 2001. Equilibrium and dynamic interfacial tension measurements at microscopic interfaces using a micropipet technique. 2. Dynamics of phospholipid monolayer formation and equilibrium tensions at water-air interface. Langmuir. 17(18):5544–5550.

10. Gaines Jr., G. L. 1966. Insoluble Monolayers at Liquid-Gas Interfaces. In 147. New York, Interscience Publishers.

11. Rugonyi, S., S. C. Biswas, and S. B. Hall. 2008. The biophysical function of pulmonary surfactant. Resp. Physiol. Neurobi. 163(1-3):244–255.

12. Zhang, H., Y. E. Wang, Q. Fan, and Y. Y. Zuo. 2011. On the low surface tension of lung surfactant. Langmuir. 27(13):8351–8358.

13. Sen, A., S.-W. Hui, M. Mosgrober-Anthony, B. A. Holm, and E. A. Egan. 1988. Localization of lipid exchange sites between bulk lung surfactants and surface monolayer: freeze fracture study. J. Coll. Interface Sci. 126(1):355–360.

14. Haller, T., P. Dietl, H. Stockner, M. Frick, N. Mair, I. Tinhofer, A. Ritsch, G. Enhorning, and G. Putz. 2004. Tracing surfactant transformation from cellular release to insertion into an air-liquid interface. Am. J. Physiol-Lung. C. 286(5):L1009–L1015.

15. Schürch, S., D. Schürch, T. Curstedt, and B. Robertson. 1994. Surface activity of lipid extract surfactant in relation to film area compression and collapse. J. Appl. Physiol. 77(2):974–986.

16. Nag, K., J. Perez-Gil, M. L. Ruano, L. A. Worthman, J. Stewart, C. Casals, and K. M. Keough. 1998. Phase transitions in films of lung surfactant at the air-water interface. Biophys. J. 74(6):2983–2995.

17. Dagan, M. P., and S. B. Hall. 2015. The Equilibrium Spreading Tension of Pulmonary Surfactant. Langmuir. 31(48):13063–13067.

18. Veldhuizen, E. J. A., J. J. Batenburg, L. M. G. van Golde, and H. P. Haagsman. 2000. The role of surfactant proteins in DPPC enrichment of surface films. Biophys. J. 79(6):3164–3171.

19. Watkins, J. C. 1968. The surface properties of pure phospholipids in relation to those of lung extracts. Biochim. Biophys. Acta. 152(2):293–306.

20. Clements, J. A. 1977. Functions of the alveolar lining. Am. Rev. Respir. Dis. 115(6 part 2):67–71.

21. Bangham, A. D., C. J. Morley, and M. C. Phillips. 1979. The physical properties of an effective lung surfactant. Biochim. Biophys. Acta. 573(3):552–556.

22. Kaganer, V. M., H. Möhwald, and P. Dutta. 1999. Structure and phase transitions in Langmuir monolayers. Rev. Mod. Phys. 71(3):779–819.

23. Veldhuizen, R., K. Nag, S. Orgeig, and F. Possmayer. 1998. The role of lipids in pulmonary surfactant. Biochim. Biophys. Acta. 1408(2-3):90–108.

24. Albrecht, O., H. Gruler, and E. Sackmann. 1981. Pressure-composition phase diagrams of cholesterol/lecithin, cholesterol/phosphatidic acid and lecithin/phosphatidic acid mixed monolayers: a Langmuir film balance study. J. Colloid. Interf. Sci. 79:319–338.

25. Crane, J. M., G. Putz, and S. B. Hall. 1999. Persistence of phase coexistence in disaturated phosphatidylcholine monolayers at high surface pressures. Biophys. J. 77(6):3134–3143.

26. Discher, B. M., K. M. Maloney, D. W. Grainger, C. A. Sousa, and S. B. Hall. 1999. Neutral lipids induce critical behavior in interfacial monolayers of pulmonary surfactant. Biochemistry. 38(1):374–383.

27. Schürch, S., R. Qanbar, H. Bachofen, and F. Possmayer. 1995. The surface-associated surfactant reservoir in the alveolar lining. Biol. Neonate. 67 (Suppl. 1)(61):61–76.

28. Schurch, S., H. Bachofen, and F. Possmayer. 2001. Surface activity in situ, in vivo, and in the captive bubble surfactometer. Comp. Biochem. Physiol. A Mol. Integr. Physiol. 129(1):195–207.

29. Follows, D., F. Tiberg, R. K. Thomas, and M. Larsson. 2007. Multilayers at the surface of solutions of exogenous lung surfactant: direct observation by neutron reflection. Biochim. Biophys. Acta. 1768(2):228–235.

30. Ueda, S., N. Ishii, S. Matsumoto, K. Hayashi, and M. Okayasu. 1983. Ultrastructural studies on surface lining layer of the lungs. Part II. J. Jpn. Med. Soc. Biol. Interface. 14:24–46.

31. Hills, B. A. 1988. The Biology of Surfactant. In 222-235. Cambridge; New York, Cambridge University Press. 222–235.

32. Schürch, S., F. H. Green, and H. Bachofen. 1998. Formation and structure of surface films: captive bubble surfactometry. Biochim. Biophys. Acta. 1408(2-3):180–202.

33. Notter, R. H., J. N. Finkelstein, and R. D. Taubold. 1983. Comparative adsorption of natural lung surfactant, extracted phospholipids, and artificial phospholipid mixtures to the air-water interface. Chem. Phys. Lipids. 33(1):67–80.

34. Bligh, E. G., and W. J. Dyer. 1959. A rapid method of total lipid extraction and purification. Can. J. Biochem. Physiol. 37(8):911–917.

35. Borie, B. 1965. X-Ray Diffraction in Crystals, Imperfect Crystals, and Amorphous Bodies. J. Am. Chem. Soc. 87(1):140–141.

36. McGovern, I. T., D. Norman, and R. H. Williams. 1987. Surface science with synchrotron radiation. Handbook on Synchrotron Radiation. G. V. Marr, editor. Elsevier, Amsterdam, pp. 467–539.

37. Als-Nielsen, J., D. Jacquemain, K. Kjaer, F. Leveiller, M. Lahav, and L. Leiserowitz. 1994. Principles and applications of grazing incidence X-ray and neutron scattering from ordered molecular monolayers at the air-water interface. Phys. Rep. 246(5):251–313.

38. Jensen, T. R., and K. Kjaer. 2001. Structural properties and interactions of then films at the air-liquid interface explored with syndhrotron X-ray scattering. vol. 11. Novel Methods to Study Interfacial Layers. D. Möbius and R. Miller, editors. Elsevier, Amsterdam; New York, pp. 205–254.

39. Tanaka, M., M. F. Schneider, and G. Brezesinski. 2003. In-plane structures of synthetic oligolactose lipid monolayers - impact of saccharide chain length. ChemPhysChem. 4(12):1316–1322.

40. Danauskas, S. M., D. X. Li, M. Meron, B. H. Lin, and K. Y. C. Lee. 2008. Stochastic fitting of specular X-ray reflectivity data using StochFit. J. Appl. Crystallogr. 41:1187–1193.

41. Andreev, K., M. W. Martynowycz, M. L. Huang, I. Kuzmenko, W. Bu, K. Kirshenbaum, and D. Gidalevitz. 2018. Hydrophobic interactions modulate antimicrobial peptoid selectivity towards anionic lipid membranes. Biochim. Biophys. Acta. 1860(6):1414–1423.

42. Andreev, K., C. Bianchi, J. S. Laursen, L. Citterio, L. Hein-Kristensen, L. Gram, I. Kuzmenko, C. A. Olsen, and D. Gidalevitz. 2014. Guanidino groups greatly enhance the action of antimicrobial peptidomimetics against bacterial cytoplasmic membranes. Biochim. Biophys. Acta. 1838(10):2492–2502.

43. Andreev, K., M. W. Martynowycz, A. Ivankin, M. L. Huang, I. Kuzmenko, M. Meron, B. Lin, K. Kirshenbaum, and D. Gidalevitz. 2016. Cyclization Improves Membrane Permeation by Antimicrobial Peptoids. Langmuir. 32(48):12905–12913.

44. Andreev, K., M. W. Martynowycz, and D. Gidalevitz. 2019. Peptoid drug discovery and optimization via surface X-ray scattering. Biopolymers. 110(6):e23274.

45. Nobre, T. M., M. W. Martynowycz, K. Andreev, I. Kuzmenko, H. Nikaido, and D. Gidalevitz. 2015. Modification of Salmonella Lipopolysaccharides Prevents the Outer Membrane Penetration of Novobiocin. Biophys. J. 109(12):2537–2545.

46. Martynowycz, M. W., A. Rice, K. Andreev, T. M. Nobre, I. Kuzmenko, J. Wereszczynski, and D. Gidalevitz. 2019. Salmonella membrane structural remodeling increases resistance to antimicrobial peptide LL-37. ACS Infect. Dis. 5(7):1214–1222.

47. Martynowycz, M. W., B. Hu, I. Kuzmenko, W. Bu, A. Hock, and D. Gidalevitz. 2016. Monomolecular siloxane film as a model of single site catalysts. J. Am. Chem. Soc. 138(38):12432–12439.

48. Clements, J. A. 1957. Surface tension of lung extracts. P. Soc. Exp. Biol. Med. 95(1):170–172.

49. Ivankin, A., I. Kuzmenko, and D. Gidalevitz. 2010. Cholesterol-phospholipid interactions: new insights from surface x-ray scattering data. Phys. Rev. Lett. 104(10):108101.

50. Discher, B. M., K. M. Maloney, D. W. Grainger, and S. B. Hall. 2002. Effect of neutral lipids on coexisting phases in monolayers of pulmonary surfactant. Biophys. Chem. 101:333–345.

51. Neville, F., M. Cahuzac, O. Konovalov, Y. Ishitsuka, K. Y. C. Lee, I. Kuzmenko, G. M. Kale, and D. Gidalevitz. 2006. Lipid headgroup discrimination by antimicrobial peptide LL-37: Insight into mechanism of action. Biophys. J. 90(4):1275–1287.

52. Gidalevitz, D., Y. Ishitsuka, A. S. Muresan, O. Konovalov, A. J. Waring, R. I. Lehrer, and K. Y. C. Lee. 2003. Interaction of antimicrobial peptide protegrin with biomembranes. Proc. Natl. Acad. Sci. USA. 100(11):6302–6307.

53. Wu, G., J. Majewski, C. Ege, K. Kjaer, M. J. Weygand, and K. Y. C. Lee. 2005. Interaction between lipid monolayers and poloxamer 188: An X-ray reflectivity and diffraction study. Biophys. J. 89(5):3159–3173.

54. Marangoni, M. N., M. W. Martynowycz, I. Kuzmenko, D. Braun, P. E. Polak, G. Weinberg, I. Rubinstein, D. Gidalevitz, and D. L. Feinstein. 2016. Membrane Cholesterol Modulates Superwarfarin Toxicity. Biophys. J. 110(8):1777–1788.

55. Bermel, M. S., J. T. McBride, and R. H. Notter. 1984. Lavaged excised rat lungs as a model of surfactant deficiency. Lung. 162(2):99–113.

56. Hall, S. B., A. R. Venkitaraman, J. A. Whitsett, B. A. Holm, and R. H. Notter. 1992. Importance of hydrophobic apoproteins as constituents of clinical exogenous surfactants. Am. Rev. Respir. Dis. 145(1):24–30.

57. Putz, G., J. Goerke, and J. A. Clements. 1994. Surface activity of rabbit pulmonary surfactant subfractions at different concentrations in a captive bubble. J. Appl. Physiol. 77(2):597–605.

58. Veldhuizen, R., K. Nag, S. Orgeig, and F. Possmayer. 1998. The role of lipids in pulmonary surfactant. Biochim. Biophys. Acta. 1408(2-3):90–108.

59. Discher, B. M., W. R. Schief, V. Vogel, and S. B. Hall. 1999. Phase separation in monolayers of pulmonary surfactant phospholipids at the air-water interface: Composition and structure. Biophys. J. 77(4):2051–2061.

60. Discher, B. M., K. M. Maloney, W. R. Schief, Jr., D. W. Grainger, V. Vogel, and S. B. Hall. 1996. Lateral phase separation in interfacial films of pulmonary surfactant. Biophys. J. 71(5):2583–2590.

61. Mansour, H. M., and G. Zografi. 2007. Relationships between equilibrium spreading pressure and phase equilibria of phospholipid bilayers and monolayers at the air-water interface. Langmuir. 23(7):3809–3819.

62. Patlak, C. S., and N. L. Gershfeld. 1967. A theoretical treatmnt for the kinetics of monolayer desorption from interfaces. J. Colloid. Interf. Sci. 25(4):503–513.

63. Andersson, J. M., C. Grey, M. Larsson, T. M. Ferreira, and E. Sparr. 2017. Effect of cholesterol on the molecular structure and transitions in a clinical-grade lung surfactant extract. Proc. Natl. Acad. Sci. USA. 114(18):E3592–E3601.

64. Larsson, M., K. Larsson, T. Nylander, and P. Wollmer. 2003. The bilayer melting transition in lung surfactant bilayers: the role of cholesterol. Eur. Biophys. J. 31(8):633–636.

65. Larsson, M., T. Nylander, K. M. Keough, and K. Nag. 2006. An X-ray diffraction study of alterations in bovine lung surfactant bilayer structures induced by albumin. Chem. Phys. Lipids. 144(2):137–145.

66. Rapp, B., and H. Gruler. 1990. Phase transitions in thin smectic films at the air-water interface. Phys. Rev. A. 42(4):2215–2218.

67. McConnell, H. M., and M. Vrljic. 2003. Liquid-liquid immiscibility in membranes. Annu. Rev. Bioph. Biom. 32:469–492.

68. Colacicco, G., and M. K. Basu. 1977. Effects of cholesterol and cholesteryl ester on dynamic surface tension of dipalmitoyl lecithin. J. Colloid. Interf. Sci. 61(3):516–518.

69. Notter, R. H., S. A. Tabak, and R. D. Mavis. 1980. Surface properties of binary mixtures of some pulmonary surfactant components. J. Lipid Res. 21(1):10–22.

70. Hildebran, J. N., J. Goerke, and J. A. Clements. 1979. Pulmonary surface film stability and composition. J. Appl. Physiol. 47(3):604–611.

